# Multi-Drug-Resistance in Drug-Naive and Drug-Exposed Ovarian Cancer Cell Lines Responds Differently to Cell Culture Dimensionality

**DOI:** 10.1101/2020.07.12.199125

**Authors:** Vasilij Koshkin, Mariana Bleker de Oliveira, Chun Peng, Laurie E. Ailles, Geoffrey Liu, Allan Covens, Sergey N. Krylov

## Abstract

Does cell clustering influence intrinsic and acquired multi-drug resistance (MDR) differently? To address this question, we studied cultured monolayers (representing individual cells) and cultured spheroids (representing clusters) formed by drug-naïve (intrinsic MDR) and drug-exposed (acquired MDR) lines of ovarian cancer A2780 cells by cytometry of reaction rate constant (CRRC). MDR efflux was characterized by accurate and robust “cell number *vs*. MDR efflux rate constant (*k*_MDR_)” histograms. Both drug-naïve and drug-exposed monolayer cells presented unimodal histograms; the histogram of drug-exposed cells was shifted towards higher *k*_MDR_ value suggesting greater MDR activity. Spheroids of drug-naïve cells presented a bimodal histogram indicating the presence of two subpopulations with different MDR activity. In contrast, spheroids of drug-exposed cells presented a unimodal histogram qualitatively similar to that of the monolayers of drug-exposed cells but with a moderate shift towards greater MDR activity. The observed greater effect of cell clustering on intrinsic than on acquired MDR can help guide the development of new therapeutic strategies targeting clusters of circulating tumor cells.

## 1. Introduction

Ovarian cancer (OC) is the seventh most common cancer in women, and chemotherapy is its frontline treatment after surgery [1]. However, effect of chemotherapeutic treatment is limited by chemoresistance. Approximately 30% of patients have pre-intrinsic chemoresistance and others eventually acquire chemoresistance during continuing exposure to chemotherapeutic agents [2]. Because chemoresistance is among the main reasons preventing progress in OC treatment, there has been intensive research in this field. A promising new approach in elucidating the mechanisms of cancer chemoresistance is studying circulating tumor cells (CTCs). CTCs exist as individual cells or multicellular clusters that detach from the primary tumor and circulate in the bloodstream and give rise to systemic metastases which are seen in some OC patients [3]. Chemoresistance of CTC clusters is greater than that of individual CTCs [4]. Understanding the mechanisms of chemoresistance in CTCs in OC is important as there is a correlation between the number of CTC clusters and clinical features of OC [5].

There are several cellular processes that contribute to both intrinsic and acquired chemoresistance [6]. One of such processes is active extrusion of drugs from cells by ATP-binding cassette transporters (ABC transporters), which are membrane proteins [7]. This process has low drug specificity and is termed multi-drug resistance (MDR). Here, we use the term of MDR solely to describe the catalytic process of drug transport across the membrane (from inside to outside of the cell). Activity of MDR in OC tumor cells has been shown to correlate with clinical chemoresistance, and MDR is its likely driver [8].

The presumed roles of MDR transport and CTC clusters in the development of chemoresistance logically lead to a question: how does MDR transport activity of clusters differ from that of single cells in cases of intrinsic and acquired resistance? Addressing this question requires a cytometry technique capable of accurately measuring MDR activity and applicable to both single cells and intact clusters. Classical cytometry technics cannot be used for accurate measurements of MDR activity, making the above-posed question difficult to approach experimentally. In contrast, cytometry of reaction rate constant (CRRC) can be used for this purpose. In general, CRRC utilizes time-lapse fluorescence microscopy to measure a rate constant of a catalytic reaction in individual cells and, thus, facilitates accurate size determination for subpopulations of cells with distinct efficiencies of this reaction [9]. Time-lapse fluorescence images are used to build kinetic traces of substrate conversion into a product. A reaction rate constant is then found for every cell using a known mechanism of the reaction. Finally, a CRRC histogram that plots the frequencies *(i.e.* number of cells) for varying ranges of the rate constant value is used to accurately measure sizes of cell subpopulations with distinct reaction activities [9].

When applied to MDR transport, CRRC is used to record kinetics of fluorescent substrate extrusion from cells (Figure 1). The extrusion process is governed by the Michalis-Menten mechanism, which is characterized by two parameters: the maximum velocity, *V*_max_, and the Michaelis constant, *K*_M_. A ratio between these parameters is a first order rate constant of MDR transport, *k*_MDR_ = *V*_max_/*K*_M_, which can be easily determined from time dependence of intracellular fluorescence intensity of MDR substrate. CRRC histograms that plot number of cells *vs. k*_MDR_ ranges are robust towards variations in substrate concentration and observation time [9]; therefore, such histograms facilitate accurate determining the sizes of cell subpopulations with different activities of MDR [9].

**Figure 1.**
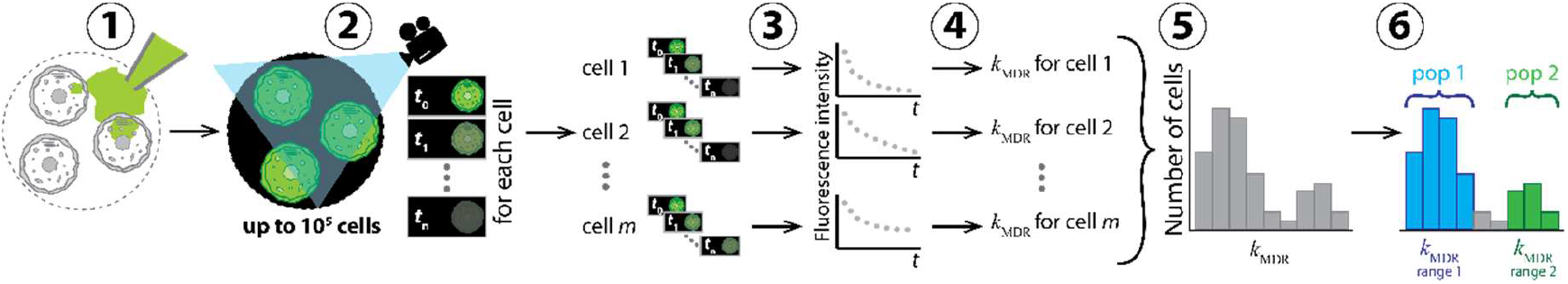
Conceptual representation of application of CRRC to MDR. The cells are loaded with a fluorescent substrate of ABC transporters which is then removed from the cell media to initiate substrate extrusion (step 1). Kinetics of decreasing fluorescence intensity is measured microscopically; sequential images of individual cells are taken over a period of time exceeding the characteristic time of the extrusion reaction (step 2). Kinetic traces of fluorescence intensity for every cell are built (step 3). Values of the reaction rate constant, *k*_MDR_, are determined for each cell (step 4). These values are used to build a CRRC histogram: “number of cells vs *k*_MDR_” (step 5). The heterogeneity of cell population with respect to MDR activity is characterized accurately using this histogram; e.g. cell subpopulations (pop) with different MDR activities are identified and quantified (step 6).

The most straightforward application of CRRC is to 2D models, such as cells cultured as monolayers or cells obtained by disintegration of cell clusters (e.g. spheroids or tissue samples) and allowed to settle on the surface [10]. However, if CRRC is based on confocal microscopy, it can also be applied to intact cell clusters [10]. Thus, CRRC is uniquely capable of accurately measuring MDR activity in both 2D and 3D models, making it suitable for addressing our question of how MDR activity of single cells differs from that of aggregated cells in (i) drug-naïve and (ii) drug-exposed tumor cells.

To address this question in the context of OC, we choose two sublines of A2780 OC cells, recently used to mimic OC circulating cells [11]. The A2780S cell subline is derived from a patient who was not exposed to chemotherapy, and, thus, this drug-naïve cell line represents intrinsic chemoresistance [12]. The A2780CP cell subline is derived from the A2780S subline and has been cultured in the presence of cisplatin to develop drug-resistance. Therefore, drug-exposed A2780CP cells represent OC tumor cells that were exposed to chemotherapy and developed acquired chemoresistance. Importantly, both cell lines can be grown as monolayers or as multicellular spheroids [13]. Therefore, we can view cultured monolayers of A2780S and A2780CP cells as models of circulating single cells, while their cultured spheroids can be considered as models of CTC clusters. Then, comparison of CRRC histograms of a cell monolayer with that of cultured spheroids can answer our question.

## 2. Results and Discussion

We grew cultured A2780S and A2780CP cells as monolayers and spheroids, loaded them with fluorescein (a fluorescent substrate of ABC transporters) and imaged fluorescein extrusion from the cells with confocal laser-scanning microscopy as described in Materials and Methods. Representative images taken at different times are shown in Figure 2.

**Figure 2.**
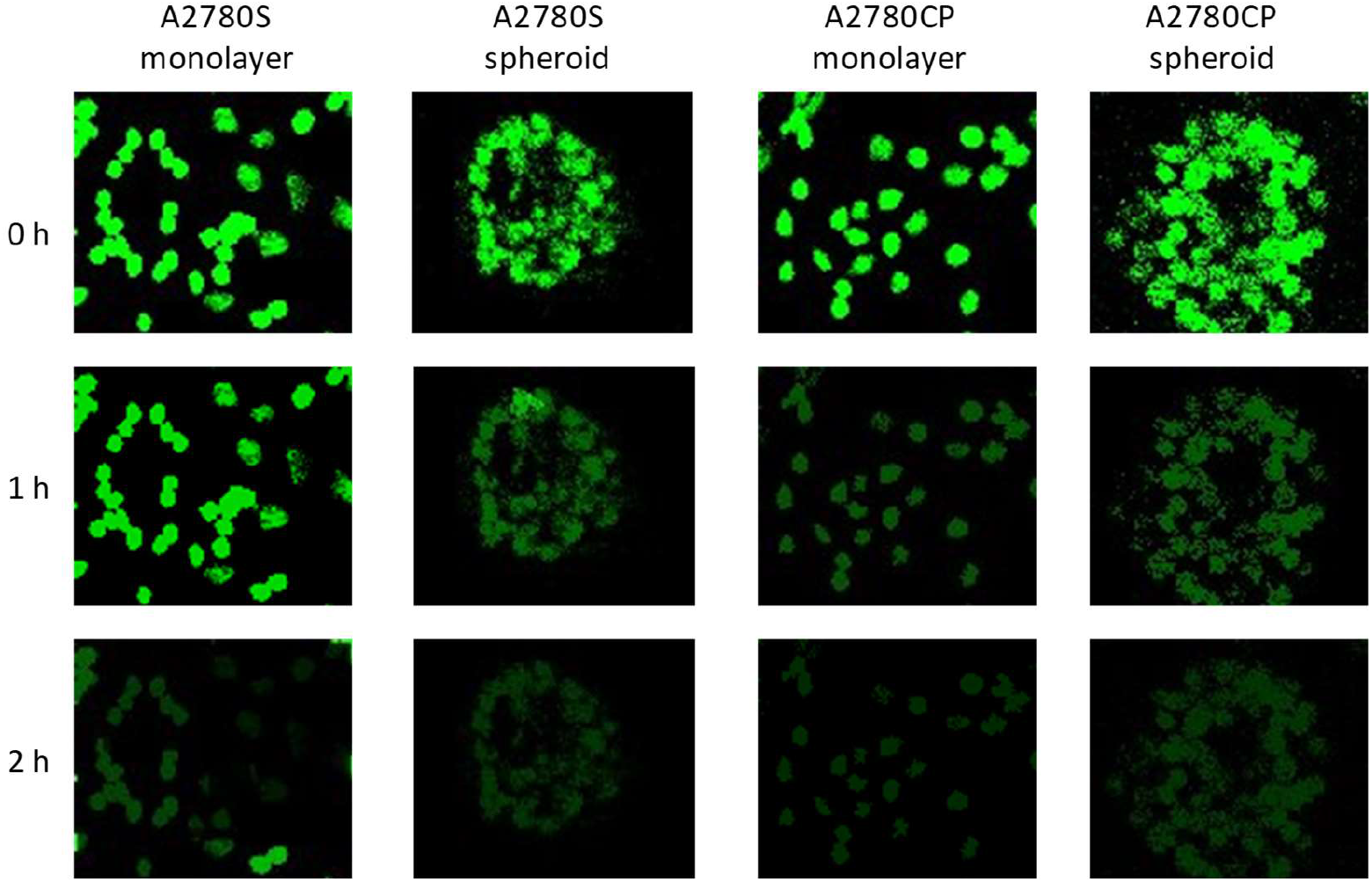
Representative images of monolayer-grown cells and spheroid-grown cells for the drug-naïve A2780S cell line and its derivative drug-exposed to cisplatin A2780CP cell line. The top, middle, and bottom images were taken at times 0, 1, and 2 h after the beginning of fluorescein extrusion.

Images of A2780S and A2780CP cells (both monolayers and spheroids) show close levels of fluorescence after loading with the same concentration of fluorescein (Figure 2, top). All cell types demonstrate also similar loss of fluorescence after completion of dye extrusion in 2 h (Figure 2, bottom). However, images taken in 1 h after the beginning of fluorescein extrusion (Figure 2, middle) suggest that A2780S monolayer cells may extrude the substrate slower than A2780S spheroid cells, as well as both types of A2780CP cells. Qualitative analysis of the images does not allow one to make any further conclusion; therefore, information in the images was a subject to CRRC kinetic analysis. We processed images from 347 cells in each of the four categories (drug-naïve single cells, drug-naïve spheroidal cells, drug-exposed single cells, and drug-exposed spheroidal cells) to determine *k*_MDR_ for each cell and plot CRRC histograms (Figure 3).

**Figure 3.**
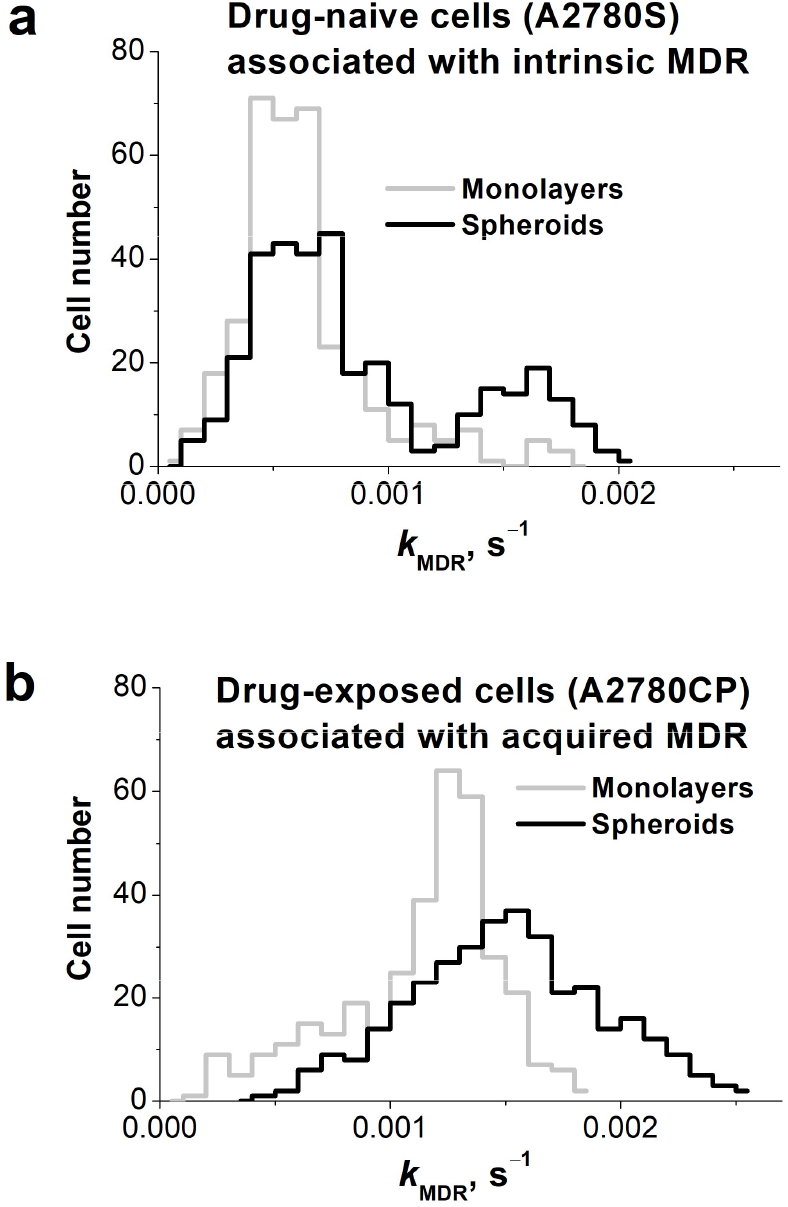
CRRC histograms of frequencies of cell subpopulations by MDR transport first order rate constant (*k*_MDR_, measured in s^−1^) in monolayer-grown cells (grey lines) and spheroid-grown cells (black lines) for drug-naïve A2780S cell line (a) and its derivative drug-exposed to cisplatin A2780CP cell line (b). Traces in panel A have been adopted with permission from Figure 4 in *Analytical Chemistry* 2020; 92, 9348–9355; Copyright 2020 American Chemical Society.

CRRC histograms of the drug-naïve A2780S cell line are shown in Figure 3a; they have been adopted from our recently published paper [10]. The monolayer histogram (grey line) was found to be unimodal, suggesting a single population of cells. The spheroid histogram (black line) revealed a bimodal distribution, suggesting two cell subpopulations: the first one is larger and has the same peak kMDR value as the monolayer cells, while the second one is smaller and has a peak *k*_MDR_ value which is almost 3 times greater. This latter subpopulation has a greater MDR capacity and its appearance is caused by cell-cell interactions in the 3D spheroids [15]. The presence of a drug-resistant subpopulation (presumably containing tumor initiating cells) in the spheroids is consistent with a notion that CTC clusters have a greater drug-resistance capacity than single CTCs. The unimodal right-skewed histogram of monolayer A2780S cells is similar to that reported earlier [9]. Its shape is consistent with the often reported asymmetric expression of MDR transporters, when the majority of cells with a basal (low) level of transporter expression form the main peak and a much smaller subpopulation of cells with the elevated level of transporter expression forms the distribution tail towards higher *k*_MDR_ values [16]. When A2780S cells are cultured as spheroids, the tail becomes a distinct peak, which corresponds to a distinct subpopulation of cells with its own elevated peak *k*_MDR_ value: the histogram becomes bimodal. This bimodality is not associated with cell position in the 3D spheroids as only the outer spheroidal cells were analysed. It is remarkable that the activation of MDR transport in spheroidal A2780S cells is not distributed equally across all spheroidal cells. Instead, the activation proceeds through enlarging the size of the subpopulation of cells with a greater MDR-transport activity. This characteristic of MDR modulation agrees well with the notion that the size and activity of the drug-resistant subpopulation determine the overall resistance of a heterogeneous cell population [17].

CRRC histograms of the cisplatin-exposed (drug-resistant) A2780CP cell line are depicted in Figure 3b. Monolayer cells (grey line) showed a unimodal distribution with the median (peak) *k*_MDR_ value exceeding that of monolayer A2780S cells (grey line in Figure 1A) approximately by a factor of 2. When cultured as spheroids, A2780CP cells also showed a unimodal distribution (black line), and the histogram of the spheroidal A2780CP cells was moderately shifted to the right with respect to that of the monolayer A2780CP cells. In addition, the peak maximum of the spheroidal A2780CP cells was at the same *k*_MDR_ position as the peak maximum of the drug-resistant subpopulation in the spheroidal A2780S cells (right-hand-side peak in the black line in Figure 1A). Thus, the drug-resistant subpopulation dominates even in monolayers formed by these cells and predictably dominates in spheroids resulting in a unimodal *k*_MDR_ distribution in the spheroid culture. Peak maximum value in the spheroid culture slightly exceeds that in monolayer (by a factor of 1.2). Lower kurtosis (indicator of distribution peakedness/flatness, −0.95 *vs* 1. 87) indicates that MDR distribution in A2780CP spheroids is more heterogeneous than in monolayers. Greater heterogeneity can be associated with the larger fraction of cells with elevated *k*_MDR_ able to survive chemotherapy and initiate tumor relapse.

The potential clinical implications of our findings are dual. First, considering the role of clustering of OC cells in intrinsic and acquired chemoresistance, our results provide a new possible explanation for the benefit of debulking surgery that has not yet been theorized; by reducing spheroids and thereby leading to less intrinsic resistance we should improve outcomes. Second, activation of MDR transport within spheroids was ascribed to activation of the HIF pathway caused either by hypoxia inside the spheroids or by cytotoxic agents [18]. However, we found MDR-transport activation not only inside the spheroids but also on their surface; moreover, this activation is observed in both drug-naïve (intrinsic MDR) and drug-exposed (acquired MDR) cells. These observations strongly suggest that there are other mechanisms of MDR-transport activation in addition to the HIF-hypoxia pathway.

To conclude, this work demonstrates unique capabilities of CRRC in studying heterogeneity of cell population with respect to MDR activity. Further, our data show that the extent of MDR-transport activation in OC cell clusters strongly depends on the previous chemotherapeutic history of spheroid-forming A2780 cells. This fact, if confirmed on primary ovarian tumor cells, will help clinicians to optimize OC treatment, since therapeutic approaches might have different outcomes for drug-naïve and drug-exposed tumors. If the observed phenomena are found in other types of cancer cells, the last conclusion can be extended to those types of cancer.

## 3. Materials and Methods

Detailed description of experimental techniques of CRRC for cell monolayers and cells in intact spheroids can be found elsewhere [10]; these techniques were followed exactly with no modification of the procedures. Briefly, monolayers and spheroids were cultured in DMEM medium supplemented with bovine serum and antibiotics under standard cell culture conditions. The culturing of small multicellular spheroids was based on the liquid overlay approach,^14^ adapted for ovarian cells [19]. Cells were placed into wells coated with agarose to prevent adhesion and allowed to form spheroids for 2–3 days. For time-lapse imaging, spheroids were placed onto coverslips and allowed to settle and attach to the surface for 5 h [20]. Imaging of MDR efflux was performed with a FV300 confocal cell imager (Olympus) in the time-lapse mode with single and multiple optical sections taken for monolayers and spheroids, respectively. Cells were loaded with a fluorescent MDR substrate (fluorescein) and allowed to extrude it. Kinetics of substrate extrusion was monitored by measuring intracellular fluorescence intensity over time to determine *k*_MDR_ for individual cells, and, importantly, only outer spheroidal cells were taken into consideration. Finally, kinetic cytometry histograms were plotted to compare MDR activity (*k*_MDR_) of monolayers and spheroids in A2780S and A2780CP cell lines (Figure 3).

## Supporting information

Key to supplementary files

## Supplementary Materials

Supplementary materials can be found at www.mdpi.com/xxx/s1.

## Author Contributions

Conceptualization, V.K., C.P., L.E.A., G.L., A.C. and S.N.K.; methodology development, V.K. and S.N.K.; performing experiments, V.K. and M.B.O.; data analysis, V.K., M.B.O and S.N.K; writing — original draft preparation, V.K.; writing — review and editing, C.P., L.E.A., G.L., A.C. and S.N.K.; supervision, S.N.K.; project administration, S.N.K.; funding acquisition, S.N.K. All authors have read and agreed to the published version of the manuscript.

## Acknowledgments

The authors thank Dr. Sven Kochmann for technical help.

## Abbreviations

ABC: ATP-binding cassette
CRRC: Cytometry of reaction rate constant
CTC: Circulating tumor cells
MDR: Multi-drug resistance
OC: Ovarian cancer

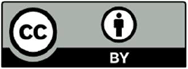 © 2020 by the authors. Submitted for possible open access publication under the terms and conditions of the Creative Commons Attribution (CC BY) license (http://creativecommons.org/licenses/by/4.0/).

