## Supplementary material for "Multi-Drug-Resistance in Drug-Naive and Drug-Exposed Ovarian Cancer Cell Lines Responds Differently to Cell Culture Dimensionality": Key to supplementary files

**Table of Contents**

| Section | Page |
|---|---|
| Supporting files description. . . . . . . . . . . . . . . . . . . . . . . . . . . . . . . . . . . . . . . . . . . . . . . . . . . . . . . . . . . . | S1 |

**Supporting Files**

| File Name | Description |
|---|---|
| ImageStacks.zip | File contains image stacks of monolayers formed by the drug-naïve (T stacks 1, 2) and drug-exposed cells (T stacks 3, 4), as well as image stacks of spheroids formed by the drug-naive (ZT stacks 1–6) and drug-exposed (ZT stacks 7-11) cells. Image stacks were used for extraction of kinetic traces utilized subsequently for the building kinetic cytometry histograms. Different T stacks and different ZT stack correspond to experiments performed on different days with new samples.<br>Each T stack contains 60 images spaced 3 min in time; recording one image took 1.69 s. ZT stacks contain different numbers of images as spheroids varied in size and required different numbers of horizontal sections with different Z positions for every time point. Accordingly ZT stacks 1–11 contain for every time point 8, 10, 7, 6, 9, 5, 8, 6, 12, 12 and 10 images, respectively. The numbers of time points in ZT stacks 1–11 are 13, 14, 8, 10, 10, 10, 11, 10, 8, 7 and 12, respectively. Totals numbers of images in ZT stacks 1–11 are 104, 140, 56, 60, 90, 50, 88, 60, 96, 84 and 120, respectively. Completion of single Z stack took approximately 10 s and one Z stack was done every 3 min. The start of every next cycle of images with varying Z can be identified by taking into account the number of images in a Z stack. For example, for ZT stack 1, a Z stack for every time point contain 8 images, accordingly the first Z stack will start with image 1, the second Z stack will start with image 9 and so on. Similarly, for ZT stack 2, a Z stack for every time point contain 10 images, accordingly the first Z stack will start with image 1, the second Z stack will start with image 11 and so on. |
| KineticTraces.zip | Kinetic traces of individual spheroid and monolayer cells obtained from time dependence of fluorescence intensity calculated from the image stacks. Column A(X) shows image number in a sequence of consecutive images separated by 3 min; accordingly, this column can be considered as time if the number is multiplied by 3 min. Each column to the right of column A(X) contains data on time dependence of relative fluorescence intensity from a single cell. |
| KineticHistograms.zip | Kinetic cytometry histograms: $k_{MDR}$ distributions in populations of spheroid and monolayer cells. The number of cells in each histogram is 347. |
